# Gut microbiota and phytoestrogen-associated infertility in southern white rhinoceros

**DOI:** 10.1101/451757

**Authors:** Candace L. Williams, Alexis R. Ybarra, Ashley N. Meredith, Barbara S. Durrant, Christopher W. Tubbs

## Abstract

With recent poaching of southern white rhinoceros (*Ceratotherium simum simum*; SWR) reaching record levels, the need for a robust assurance population is urgent. However, the global captive SWR population is not currently self-sustaining due to the reproductive failure of captive-born females. Dietary phytoestrogens have been proposed to play a role in this phenomenon, and recent work has demonstrated a negative relationship between diet estrogenicity and fertility of captive-born female SWR. To further examine this relationship, we compared gut microbial communities, fecal phytoestrogens, and fertility of SWR to another rhinoceros species–the greater one-horned rhinoceros (*Rhinoceros unicornis*; GOHR), which consumes a similar diet but exhibits high levels of fertility in captivity. Using 16S rRNA amplicon sequencing and mass spectrometry, we identified a species-specific fecal microbiota and three dominant fecal phytoestrogen profiles. These profiles exhibited varying levels of estrogenicity when tested in an *in vitro* estrogen receptor activation assay for both rhinoceros species, with profiles dominated by the microbial metabolite, equol, stimulating the highest levels of receptor activation. Finally, we found that SWR fertility varies significantly with respect to phytoestrogen profile, but also with the abundance of several bacterial taxa and microbially-derived phytoestrogen metabolites. Taken together, these data suggest that in addition to species differences in estrogen receptor sensitivity to phytoestrogens, reproductive outcomes may be driven by gut microbiota’s transformation of dietary phytoestrogens in captive SWR females.

## Background

The southern white rhinoceros (SWR; *Ceratotherium simum simum*) has returned from the brink of extinction through extensive *in situ* and *ex situ* conservation efforts, with wild populations increasing from approximately 100 to 20,000 over the last century (1). However, wild SWR now face an uncertain future due to the recent dramatic increase in poaching (2). An additional challenge facing the species is the reproductive failure of the once robust *ex situ* assurance populations (3,4). Together, poaching, long gestational length (∼16 months) and inter-calving interval (∼ 2.5 years) (5), and captive infertility (3,4) have rendered both wild and captive populations no longer self-sustaining. Without any change in poaching rates, wild SWR populations will likely face the threat of extinction within the next two decades (6).

Previous work has implicated captive diets in the reproductive failure of captive SWR (4,7). In the wild, SWR are pure grazers, consuming up to ∼40 kg/day of various grasses (8,9). In contrast, diets in managed settings typically contain phytoestrogen-rich legume hays and soy-and alfalfa-based concentrated feeds (4). A survey of nine SWR-breeding institutions demonstrated that diet estrogenicity was strongly associated with the amount of soy and/or alfalfa-based pellets fed. Moreover, female SWR born at institutions feeding highly estrogenic diets exhibit lower fertility than female SWR born at institutions feeding low phytoestrogen diets (4).

Due to their structural similarity to endogenous estrogens, phytoestrogens may interact with estrogen receptors (ERs) and disrupt normal endocrine function, reproduction, and development (10–13). Previously, we showed that SWR ERs exhibit higher maximal activation by phytoestrogens than ERs of the greater one-horned rhinoceros (*Rhinoceros unicornis*; GOHR) (5). Both species consume similar high-phytoestrogen diets in captivity, but GOHR do not exhibit the decrease in fertility observed in SWRs. These data suggest that at the receptor level, SWR are particularly vulnerable to the deleterious effects of phytoestrogen exposure. Whether SWR possess additional species-specific characteristics that predispose them to phytoestrogen sensitivity remains unclear.

Due to the limitations of collecting biological samples from a threatened megafaunal species, little is understood about the specific physiological consequences of SWR consuming estrogenic diets. Altered endocrine and reproductive function by phytoestrogen exposure has been described in humans, rodents, and livestock species (11,13,14). Many of these effects, including reproductive tract pathologies, erratic or absent luteal activity, and reduced fertility, parallel findings in captive female SWR (15–17). However, the potential role of phytoestrogens in the onset of these pathologies has not been investigated. In other species, the physiological outcomes of phytoestrogen exposure are profoundly affected by transformation of parent compounds following consumption. For example, in ewes, reproductive pathologies and infertility develop following consumption of diets high in the isoflavone daidzein, but it is equol, a daidzein metabolite, that is thought to be the driver of this effect (10). Equol production relies exclusively on microbial transformation, and several other phytoestrogens are metabolized by members of the gastrointestinal tract microbiota to produce metabolites that vary in estrogenicity (18–21). Coumestrol, a compound from another class of phytoestrogens, the coumestans, also has been associated with sheep infertility (9), but to date, the microbial metabolism of coumestans has not been explored. Whether gut microbiota may play a similar role in SWR responses to dietary phytoestrogens is unclear.

The relationship between animals and their associated microbes is important, as microbiota are essential for many biological processes within their hosts (22). However, an understanding of how interactions between phytoestrogens and resident gut microbiota may affect fertility is lacking for any vertebrate species. Given what is known about bioactivation of phytoestrogens by gut microbiota in other mammalian species (23) and the strong link between dietary phytoestrogens and reproductive failures in rhinoceros (4), an investigation into phytoestrogen metabolism by rhinoceros gut microbiota is warranted. To examine these interactions, we characterized SWR and GOHR fecal microbiota as a proxy for gut microbiota. In addition, we compared fecal phytoestrogen composition and metabolite profile estrogenicity, using mass spectrometry and ER activation assays, respectively, between the two species. By sampling separately housed, but similarly managed SWR and GOHR females from the same institution, we sought to reduce variation by eliminating known drivers of gut microbiota composition, such as diet and geographic location (24–26), to better identify species differences. Finally, we used historical breeding records to examine the relationships between specific microbial taxa, phytoestrogen metabolites, and SWR reproductive success. With these data, we shed light on the role microbiota may play in captive SWR infertility with the aim to develop techniques to support and increase this species’ assurance population.

## Results

### Composition of fecal microbiota, but not phytoestrogens, differ by species

Sequencing of 16S rRNA fecal samples (SWR: *n* = 42; GOHR: *n* = 16; Table S1) from eight individual rhinoceros (SWR: *n* = 6; GOHR: *n* = 2; Table S1) revealed that GOHR samples had significantly higher inter-sample diversity compared to SWR despite SWRs having a higher number of unique, low (>1 %) relative abundance operational taxonomic units (OTUs) overall (Table S2). Significant differences in fecal community structure and composition between rhino species were also observed at the phylum, family, and OTU level using permutational analysis of variance (PERMANOVA) and accounting for relative abundances using weighted UniFrac (all *P* < 0.001). A difference in microbial communities was also observed by nonmetric multidimensional scaling (nMDS; Fig. 1, inset). Members of four phyla were found to significantly contribute to variation (Fig. 1), with the relative abundance of the Bacteroidetes (SWR: 55 ± 1.1 %; GOHR: 30 ± 1.8 %) and the Firmicutes (SWR: 33 ± 1.2 %; GOHR: 55 ± 2.2 %) found to significantly differ with respect to rhino species (Welch’s *t-test,* both *P* < 0.001; Fig. 1). Several members of these phyla were also found to be significantly different at both the family and OTU level, with six families and eleven OTUs contributing to these significant differences (Fig. 1).

**Figure 1.**
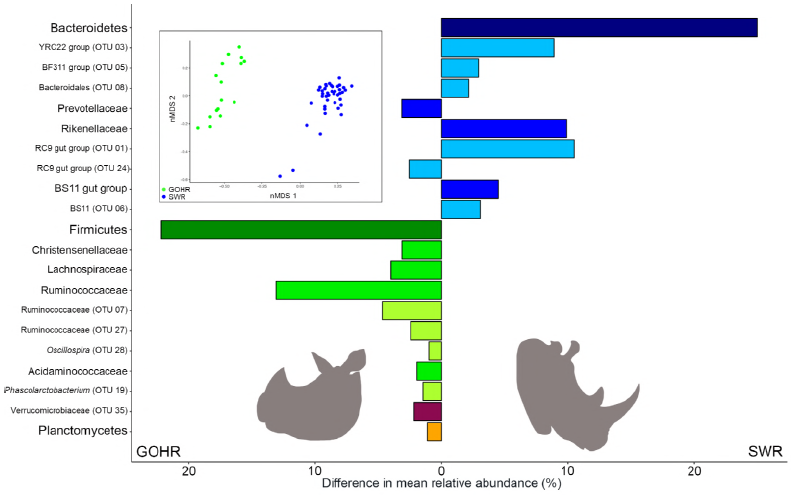
Differences in fecal microbiota between southern white rhinoceros (SWR) and greater one-horned rhinoceros (GOHR). Nonmetric multidimensional scaling (nMDS, inset) analysis displaying differences in microbiota observed by 16S rRNA amplicon sequencing based on Bray-Curtis distances (PERMANOVA, *P* < 0.001; stress: 0.13). Differences in mean relative abundance of bacterial taxa found to significantly contribute to variation between rhinoceros species (SIMPER ≥ 2.0 %; Welch’s *t-test, P* < 0.05) are organized by color, with all members of a particular phylum sharing a similar color, with intensity decreasing from phylum to family to OTU level

The observed differences in microbial community are likely related to the different foraging strategies exhibited by the two species. All individuals in this study live in large exhibits where they are provided diet of soy and alfalfa based pellets supplemented with either grasses and browse. SWR, which in the wild are grazers, consume additional hay and fresh grasses (8,9). In contrast, GOHR, a predominantly browsing species, consume a more varied diet that includes fruits and leaves (27). This difference in foraging may be driving species differences in gut microbiota, as observed in other closely related species (28). Nevertheless, both species are herbivorous and their gut microbiota are similar in that the dominant microorganisms present in both species are those capable of fiber degradation and therefore fulfill similar functional niches (29).

Despite species differences in microbial communities, neither overall structure nor composition of detected phytoestrogen analytes varied significantly between SWR and GOHR (Fig. 2A, PERMANOVA, *P* > 0.05). However, species differences were observed at the individual analyte level. Concentrations of equol (EQ), enterolactone (EL), methoxycoumestrol (MOC), and coumestrol (CO) were significantly higher in the GOHR (Fig. 2B-D, Table S3). Several phytoestrogens were detected exclusively in the diet, the isoflavones, formononetin (FM) and genistein (GN) (Fig. 2B), whereas microbially-derived metabolites EQ, 4’-ethylphenol (PEP), EL, and enterodiol (ED) were detected only in feces (Fig. 2BC). Two other phytoestrogens, biochanin-A and *o*-demethylangolesin, were not detected in any sample type (both, < 65 ppb). In general, there was an overall trend for excreted quantities of phytoestrogens and metabolites to be higher in GOHR compared to SWR (Fig. 2B-D).

**Figure 2.**
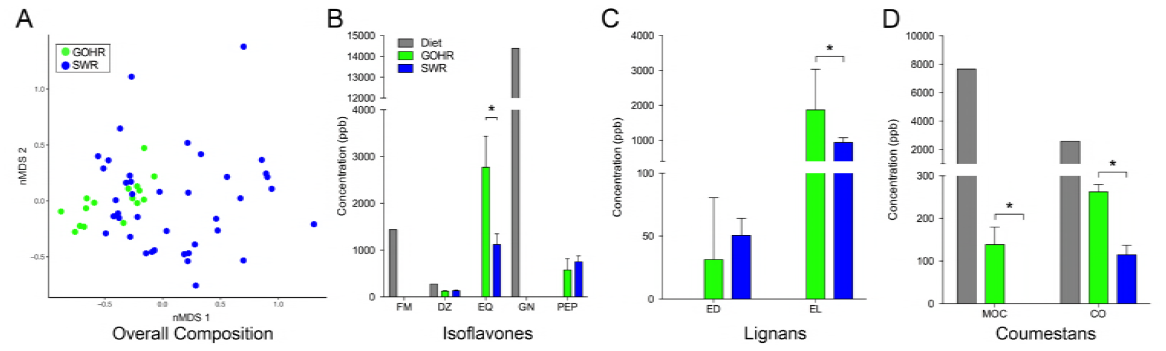
Comparison of fecal phytoestrogen composition between southern white rhinoceros (SWR) and greater one-horned rhinoceros (GOHR) A) Nonmetric multidimensional scaling (nMDS) analysis displaying overall composition of fecal phytoestrogens detected by mass spectrometry based on Bray-Curtis distances (PERMANOVA, *P* > 0.05; stress: 0.13). Mean ± SE analyte concentrations in parts per billion (ppb) of B) isoflavones, C) lignans, and D) coumestans for both SWR and GOHR and their diet. *Significantly different concentrations of fecal analytes (Welch’s *t*-test, *P* < 0.05). FM: formononetin, DZ: daidzein, EQ: equol, GN: genistein, PEP: 4’ethylphenol, ED: enterodiol, EL: enterolactone, MOC: methoxycoumestrol, CO: coumestrol.

The relative abundances of specific OTUs provide some insight into the observed phytoestrogen and metabolite concentrations described above. Overall, 77 OTUs were found to significantly correlate with phytoestrogen concentration (Fig. 3A), which were overall significantly more abundant in SWR compared to GOHR (Welch’s *t-test, P* < 0.0001; Fig. 3B). Interestingly, the 27 OTUs negatively correlated with phytoestrogen metabolite concentration are nearly four times more abundant in SWR, while the 41 positively correlated OTUs are approximately 5-fold less abundant in SWR (both, Welch’s *t-test, P* < 0.0001; Fig. 3CD). Taken together, these findings provide a plausible explanation for why there was an overall trend of lower concentrations of individual phytoestrogen analytes in SWR samples.

**Figure 3.**
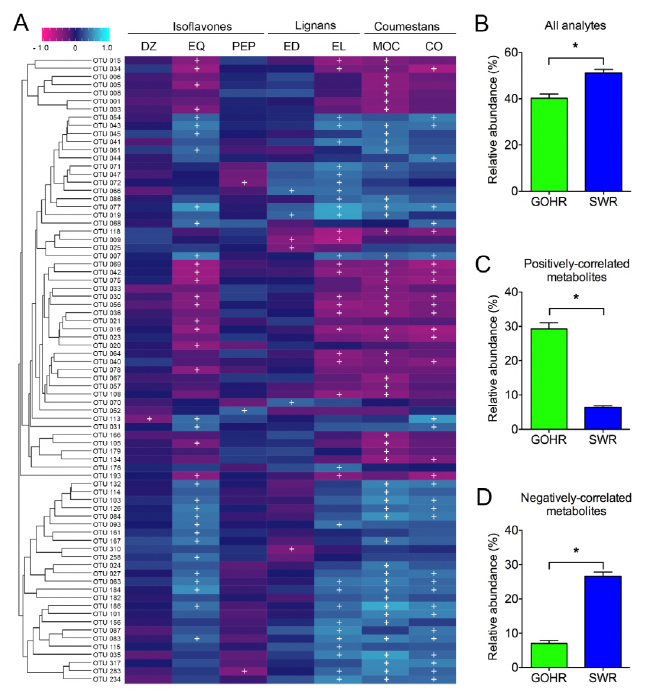
Relative abundance of OTUs and phytoestrogen concentrations significantly correlate. A) Heatmap depicting significant correlations between phytoestrogen analytes and microbiota (≥ 1.0 % relative abundance) using Spearman correlation method with FDR correction (^+^Significance at *P* < 0.05). Dendogram displays OTUs that commonly co-occur by hierarchical clustering (Bray-Curtis). Species differences in mean ± SE relative abundance of observed OTUs correlating to B) phytoestrogen analytes, C) positively correlated metabolites, and for D) negatively correlated metabolites. * Welch’s *t-test*, *P* < 0.05. DZ: daidzein, EQ: equol, PEP: 4’ethylphenol, ED: enterodiol, EL: enterolactone, MOC: methoxycoumestrol, CO: coumestrol, GOHR: greater one-horned rhinoceros, SWR: southern white rhinoceros.

One possible explanation for similarity in phytoestrogen composition between SWR and GOHR is that the OTUs positively associated with phytoestrogen concentrations belong to taxa of known fiber degraders. Many of these taxa, like members of the Bacteroidetes (Rikenellaceae) in SWR, and the Firmicutes (Ruminococcaceae and Lachnospiraceae) in GOHR accomplish this via β-glucosidase activity, as this enzyme also catalyzes early steps of phytoestrogen transformation (23). Thus, the lack of species differences in phytoestrogen composition may be driven by the overall functional similarity of the two species’ gut microbial communities.

### Three distinct phytoestrogen profiles examined

With no clear species difference in metabolite composition, hierarchical clustering was used to group similar fecal samples from both species of rhinoceros according to their phytoestrogen composition. This approach identified three distinct phytoestrogen profiles representing the most commonly observed fecal metabolite profiles in individual samples from both SWR and GOHR (Fig. 4A-C, Fig. S1, Fig. S1D). For the two most similar profiles, the moderately estrogenic EQ was the dominant metabolite produced, followed by the weakly estrogenic EL (Profile B & C; Fig. 4BC, Table S3, Fig. S1). However, total phytoestrogen concentrations in Profile C were approximately twice the total concentration of phytoestrogens detected Profile B (8,884 ± 970 ppb and 4,254 ± 315 ppb, respectively; Fig. 4BC, Fig. S1). A third, less similar profile was also identified, in which the dominant metabolite was EL (Profile A; Fig. 4A, Table S3). The total concentration of phytoestrogens in this profile was significantly lower (1,510 ± 229 ppb) (Fig. 4A, Fig. S1). Despite no visual difference in the overall communities using nMDS (Fig. S1E), several bacterial taxa were found to differ significantly with respect to phytoestrogen profiles (OTU 03, YRC22; OTU 07, OTU 27 Ruminococcaceae). However, no individual OTU contributed to variation > 8.5 %, indicating that a group of microbiota, not individual OTUs, may be important in driving differences between phytoestrogen profiles.

**Figure 4.**
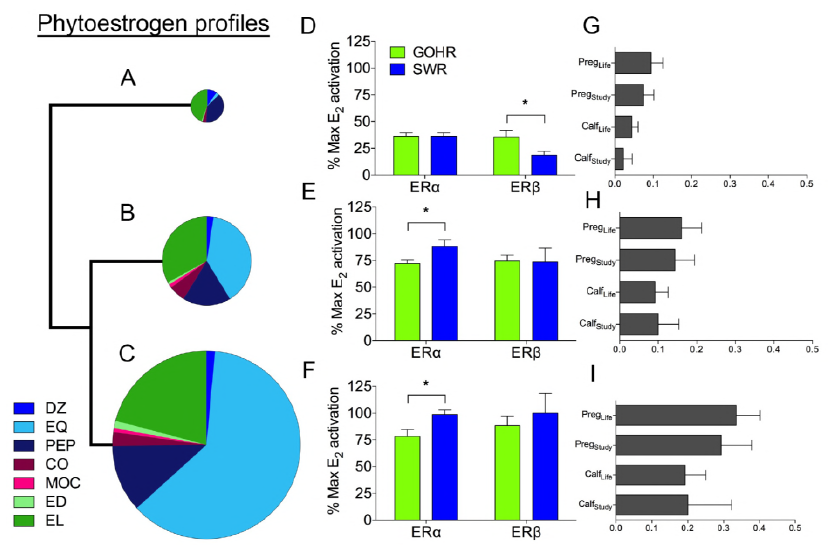
Relative estrogenicity and fertility of phytoestrogen profiles identified by hierarchical clustering. Phytoestrogen composition, as depicted by hierarchical clustering, with each profile’s size relative to total concentration detected by mass spectrometry, A) Profile A, B) Profile B, and C) Profile C. The mean ± SE activation of ERα and ERβ of both southern white rhinoceros (SWR) and greater one-horned rhinoceros (GOHR), relative to maximal activation by 17β-E_2_ by respective phytoestrogen profiles D) Profile A, E) Profile B, and F) Profile C, when tested at concentrations found *in vivo*. Differences in mean ± SE fertility measurements with respect to phytoestrogen profiles, G) Profile A, H) Profile B, and I) Profile C, *Significantly different activation (ANOVA, *P* < 0.05). DZ: daidzein, EQ: equol, PEP: 4’ethylphenol, ED: enterodiol, EL: enterolactone, MOC: methoxycoumestrol, CO: coumestrol.

To quantify the relative estrogenicity of phytoestrogen profiles (Profiles A, B, C), each observed mixture was formulated *in vitro* and tested in estrogen receptor (ER) activation assays using ERα or ERβ from SWR and GOHR (Fig. 4D-F, Fig. S1F-G), as described previously (5). All three phytoestrogen profiles activated SWR and GOHR ERs (Fig. 4D-F, Fig. S1F-G), with Profile C, the most potent agonist for both SWR ERs reaching maximal activation relative to 17β-estradiol (E_2_) (Fig. 4F, Fig. Fig. S1F-G, Table S4). Similarly, Profile B stimulated maximal activation of SWR ERα, and near maximal activation of SWR ERβ relative to E_2_, despite having less than half the total concentration of analytes of Profile C (Fig. 4E, Fig. Fig. S1F-G, Table S4). We attribute this high activation primarily to EQ, which is a dominant metabolite in both profiles and a known potent agonist to rhino ERs (5). However, it is interesting that activation of SWR ERα by Profiles B & C was significantly greater than that of GOHR ERα (Table S4), as previous work has shown GOHR ERα to be more sensitive to EQ than its SWR homologue (5). In contrast, the least potent Profile A stimulated significantly greater activation of GOHR ERβ relative to SWR ERβ (Fig. 4D, Fig. Fig. S1F-G, Table S4). This is also noteworthy, as no phytoestrogen tested in previous studies has ever been shown to be a more potent agonist of GOHR ERβ than SWRβ. What is driving these differences is unclear, as the dominant metabolites in Profile A, EL and PEP, do not appreciably bind or activate ERβs from either species, and the known agonists present in Profile A (DZ, EQ and CO) are more potent activators of SWR ERβs than those from GOHR (5). Nevertheless, this observation highlights the importance of evaluating the effects of mixtures of suspected endocrine disrupting chemicals on receptors, in addition to individual chemicals, as this method better mimics *in vivo* conditions.

### Interactions with SWR fertility explored

To assess fertility of our SWR population, the number of pregnancies achieved and/or calves born were determined for both the period of sample collection as well as for the lifetime of each of the SWR included in this study (Table S1). Pregnancies achieved (Pregnancy_study_, PS; Pregnancy_life_, PL) were confirmed via elevations in fecal progestagen levels and were included in the analysis since rhino gestation length (∼16 months) exceeded the duration of sample collection (4 months). Fertility (Calf_study_, CS; Calf_life_, CL) represents calves born per reproductive year using calculations described previously (4). When comparing phytoestrogen profiles using CS, we did not find any significant difference in mean fertility (Fig. 4G-I; Table S5). Using the PS calculation, however, we showed that individuals exhibiting Profile A had the lowest mean pregnancy rate, and those producing Profile C had the highest (Fig. 4G & I). For lifetime measures, we found a similar relationship, with PL and CL for Profile C producers being significantly greater than Profile A producers (Fig. 4G & I). Although not significantly different, SWR producing Profile B profiles tended to have higher mean fertility than individuals belonging to Profile A across all measures (Fig. 4GH).

Mean reproductive success, in terms of pregnancies achieved and calves born, was highest in individuals with the greatest concentrations of fecal metabolites (Fig. 4I; Profile C). For some of the metabolites produced, these findings parallel observations by others. For example, all measures of SWR fertility were positively correlated with production of EL (Table S5). This finding is consistent with studies in humans that have demonstrated a link between high levels of EL and increased reproductive success (31). Our previous work shows EL does not appreciably bind or activate SWR ERs and therefore possesses little endocrine disrupting potential as a xenoestrogen (7). However, the positive relationship between EQ and calf-based fertility measures is unexpected (Table S5). *In vitro*, EQ is a relatively potent agonist of both SWR ERα and ERβ (7), and in other vertebrate species EQ is cleared from the circulation less quickly than other isoflavones, increasing its bioavailability (32). This suggests that high levels of EQ production should negatively affect SWR fertility, as is well documented in other grazing species (11,12).

Also unexpected was the finding that individual SWR producing the most estrogenic profiles (Profile B & C) exhibited the highest fertility (Fig. 4HI), while SWR fertility was lowest in individuals producing profiles with the lowest overall estrogenicity (Profile A). These observations lead to several new questions. Do SWR belonging to Profile A produce novel phytoestrogen metabolites that are more estrogenic? We observe high levels of certain compounds, such as MOC and CO in feeds, but low levels are detected in feces. CO is a potent SWR ER agonist (7) and has been associated with infertility in sheep (9), but little is known about possible microbial metabolites and their relative estrogenicity. It is possible that these coumestans are converted into a novel metabolite that could be highly estrogenic to SWR. Another possible explanation for the positive association between profile estrogenicity and fertility is that the varying degrees of fecal profile estrogenicity result from differences in phytoestrogen absorption or excretion between individuals. Specifically, it could be hypothesized that elevated excretion of phytoestrogens and metabolites would reduce circulating levels, thus limiting the potential for these chemicals to cause reproductive harm. This is supported not only by our findings in individual animals, but also by our species-level observations where the more fertile GOHR generally excrete higher levels of phytoestrogens than the less fertile SWR. This does not appear to be case in in sheep and cattle, where concentrations of phytoestrogens and metabolites in excreta (i.e., urine) generally correlate to plasma levels (33–35). However, detailed studies examining the generation and clearance of phytoestrogen metabolites, and their subsequent endocrine disrupting effects on target tissues, are lacking even for relatively well-studied species. Addressing such relationships in SWR will be challenging, if not impossible. Nevertheless, the findings presented here do provide the opportunity to apply potentially innovative approaches, like using non-targeted mass spectrometry to identify novel metabolites, or using fecal EL or EQ concentrations to identify individual SWR with high reproductive potential.

Few studies have examined the interaction between mammalian fertility and gut microbiota, and defining this link is difficult. Here, we found the abundance of six OTUs to correlate to fertility measures (Table S6). Of the four OTUs that negatively correlate to fertility, three also have significant negative correlations to EL and EQ. However, OTUs positively correlated to fertility did not display significant correlations to any phytoestrogen measured in our study (Table S6), and it is unknown what role these microbiota may play. Using correlations between microbial abundance and phytoestrogen metabolites to determine microbial activity is biased, as compositional data, like presented here, do not directly correlate to microbial activity (36). Therefore, it is possible that less abundant taxa may significantly contribute to the transformation of phytoestrogens. For example, members of the Coriobacteriaceae, which have been shown to convert DZ to EQ (30), the Eubacteriaceae, which are capable of dehydroxylation of lignans to produce ED and EL (37), and the *Blautia* spp., which have displayed in both lignan and isoflavone metabolism (20, 30) are found in samples collected from both SWR and GOHR in low abundances (all, < 1.0 %). However, further work is needed to determine their contributions to phytoestrogen metabolism within the rhinoceros, including *in vitro* culture experiments to measure their microbial activity.

## Discussion

Working with threatened species, such as the two species studied here, presents its own unique set of challenges. Despite these challenges, however, our combining of parallel sequencing, mass spectrometry, and estrogen receptor activation assays, provides insight into the host-microbe relationship with fertility that, to our knowledge, is novel for any vertebrate species. Such an approach is needed to understand and apply novel application of techniques within nontraditional systems.

Our work sheds light on how microbiota may drive reproductive outcomes in SWR, but they are not the only species that may benefit from the work presented here. Among its broader application to other vertebrates, our findings may be critical for the management of SWR’s closest relative, the northern white rhinoceros (*Ceratotherium simum cottoni*, NWR), a subspecies with only two living members (38). Like SWR, NWR experience low fertility and a prevalence of reproductive pathologies in managed settings (17). As a closely related, grazing subspecies, it is likely sensitive to phytoestrogens as well. Currently, several rescue attempts are underway to prevent NWR extinction (39,40). Should these attempts to save the NWR be successful and with SWR facing a similar uncertain fate, any novel approaches to promote high fertility, such as managing microbial phytoestrogen transformation by altering microbiota through diet modifications and other therapeutic approaches, will be needed. With the information presented here, we plan to direct future work aimed at developing strategies to improve captive SWR reproduction, with the ultimate goal of alleviating their threat of extinction.

## Methods

### Study animals

Female greater one-horned rhinoceros (*n =* 2) and southern white rhinoceros (*n =* 6) used in this study were housed at the San Diego Zoo Safari Park, Escondido, CA, USA in two separate 24 ha mixed species exhibits (Table S1). All procedures were approved by San Diego Zoo Global’s Institutional Animal Care and Use Committee (#15-013).

### Sample collection

Fresh fecal samples (SWR, *n =* 42; GOHR, *n =* 16, Table S1) were collected weekly beginning September 3, 2015 through January 1, 2016, alternating weeks between SWR and GOHR. Samples were collected from animals at the same time of day using binoculars to identify individuals based on their unique horn structure. Following defecation, collection occurred between one to twenty minutes, and samples were transported on dry ice and stored at −80 °C prior to processing.

### DNA extraction

Total genomic DNA from fecal samples and negative control was extracted via mechanical disruption and hot/cold phenol extraction following Stevenson *et al.*’s protocol (2007) with the exception that 25:24:1 phenol:chloroform:isoamyl alcohol was used in place of phenol:chloroform at all steps. DNA was quantified using a Qubit Fluorometer (Invitrogen, Carlsbad, CA, USA) and stored at −20 °C following extraction.

### Library Preparation & Sequencing

Sequencing library preparation was carried out following manufacturer’s recommendations (Illumina, 2013) with some modifications. In brief, amplicon PCR targeted the V4 region of the 16S rRNA gene using a forward (V4f: TATGGTAATTGTGTGCCAGCMGCCGCGGTAA) and reverse (V4r: AGTCAGTCAGC CGGACTACHVGGGTWTCTAAT) primers in a 25-μL reaction with 1X KAPA HiFi Hot Start Ready Mix (Kapa Biosystems), 0.2 mM each primer, and 1.0 - 5.0 ng DNA (31). Amplification conditions were as follows: 95 °C for 2 min, 25 cycles of 95 °C for 20 s, 55 °C for 15 s, 72 °C for 30 s, and a final 10 min extension at 72 °C. PCR products were purified via gel extraction (Zymo Gel DNA Recovery Kit; Zymo, Irvine, CA) using a 1.0 % low melt agarose gel (National Diagnostics, Atlanta, GA) and quantified with a Qubit Fluorometer (Invitrogen). With the negative control producing no band, the expected area was excised. All samples were combined to yield an equimolar 4 nM pool. Following manufacturer’s protocol, sequencing was conducted on an Illumina MiSeq using reagent kit V2 (2 x 250 bp cycles), as described previously (Illumina, 2013). All sequences were deposited into the National Center for Biotechnological Information’s Short Read Archive under Accession Number SRP136468.

### 16S rRNA sequence analyses

Sequence analysis was carried out using mothur v.1.39.5 (43) following the MiSeq SOP (42). In brief, contigs were formed from 16S rRNA reads, and poor quality sequences were removed. Sequences were trimmed and filtered based on quality (maxambig = 0, minlength = 250, maxlength = 500). Unique sequences were aligned against the SILVA 16S rRNA gene alignment database (44) and classified with a bootstrap value cutoff of 80, and operational taxonomic units (OTUs) found with < 2 sequences in the total dataset were removed. Chimeras (chimera.uchime) and sequences identified as members of Eukaryota, Archaea, Cyanobacteria lineages, and mitochondria were also removed. Sequences were clustered into OTUs at a 97 % similarity cutoff using OptiClust (OTU table, Table S7). Negative control yielded 273 sequences, comprised of low-level cross-sample contaminants; therefore, OTUs were not removed from dataset.

Sequence coverage was assessed in mothur by rarefaction curves (Fig. S2) and Good’s coverage (45). Samples were then iteratively subsampled 10 times to 6,825 sequences per sample, and OTU abundances were calculated as whole number means across iterations. Additionally, richness and diversity were calculated for each sample. All other calculations were carried out in R using both *vegan* and *phyloseq* packages (46,47). The similarity indices Bray-Curtis (48), Jaccard (49), and weighted UniFrac (50) were used to assess differences in bacterial community, and these differences were visualized by nonmetric multidimensional scaling plots (nMDS, iters=10,000) (51). Permutational analysis of multivariate dispersions (PERMDISP2) was used to test for heterogeneity of community structure and composition between rhino species, and with unequal variances observed, data were down-sampled to create even sample sizes using the *caret* package (52) prior to permutational analysis of variance (PERMANOVA, *vegan*::adonis, SWR, *n =* 16; GOHR, *n =* 16) to determine species differences. Similarity percentages (SIMPER, *vegan*) analyses then determined the contributions from each taxonomic group to PERMANOVA reported differences. Species-related differences in individual OTUs were examined by Welch’s *t*-test (two-sided, SWR, *n =* 16; GOHR, *n =* 16). All data are expressed as the mean ± SE and considered significant if *P* < 0.05 unless otherwise stated.

### Phytoestrogen extraction and quantification

Samples collected were batched into groups of ten and accompanied by quality control samples. Phytoestrogens were extracted from fecal samples by a two-phase extraction as described previously by Palme *et al*. (2013) with few modifications. In the first phase, fecal samples were diluted ten-fold using 80 % methanol in water (Fisher Scientific), homogenized for 20 min using a Geno/Grinder^®^ at 1,000 rpm, centrifuged for 10 min at 4,000 x *g*, and the supernatant was recovered. In the second-phase, 1.0 mL of methanol extract was added to 4.0 mL diethyl ether (Fisher Scientific), 0.5 mL of 5.0% NaH_2_CO_3_ (Sigma), and 4.0 mL of water (54), inverted four times, and centrifuged for 10 min at 4000 x *g*. The ether phase was removed, evaporated at 45 °C by a nitrogen flow of 0.4 psi and resuspended in methanol. Extracts were further filtered (0.22 μm) and analyzed by liquid chromatography-coupled tandem mass spectrometry (LC/MS/MS) for all analytes with the exception of 4’-ethylphenol (PEP) which was analyzed by gas chromatography-mass selective detector (GC/MSD). Quality control samples included a blank matrix sample (grass) that was absent of phytoestrogens to assess contamination during the extraction and a matrix spiked sample which was fortified with a known concentration of phytoestrogens. The spiked matrix sample was used to determine the efficiency of the extraction for every batch; recoveries ranged between 50-150 %.

### LC/MS/MS method

Analysis was performed using an Agilent 1260 liquid chromatograph coupled to an Agilent 6430 Triple Mass Spectrometer. Chromatographic separation was performed using an Agilent Zorbax Eclipse Plus (2.1 x 50 mm id, 1.8 μm) Rapid Resolution column maintained at 40 °C. The mobile phases consisted of 5 mm ammonium formate and 0.1 % formic acid in water for the aqueous phase (A), with 5 mm ammonium formate and 0.1 % formic acid in methanol as the organic phase (B). The flow rate was held at 0.4 mL/min and gradient program was as follows: 0-0.5 min 10 % B, 0.5-3.0 min increasing to 90 % B. The ionization of phytoestrogens was performed using electrospray ionization (ESI) in positive mode with an auxiliary gas (N_2_), source temperature of 300 °C, and a gas flow rate of 12 L/min, with the exception of enterodiol which was run in negative mode. Optimized MRM conditions are listed in Table S8.

### GC/MSD method

The analysis of PEP (Indofine, CAS: 123-07-09) was performed on an Agilent 7890B gas chromatograph (GC) coupled to an Agilent 5977A mass selective detector (MSD). The GC inlet temperature was set to 280 °C run in pulsed splitless mode with an injection volume of 1 μL. The GC oven temperature was set to 80 °C and increased to 200 °C between 1 and 13 minutes at a rate of 10 °C/min. The oven temperature was then increased to 300 °C between 13 and 22 minutes at a rate of 25 °C/min for a total run time of 22 minutes. Ultra high purity helium (carrier gas) was used at a constant flow rate of 1.5 mL/min with an Agilent DB-5MS UI (30 m x 0.250 mm) 0.25 μm analytical column. PEP was analyzed using an electron ionization (EI) source with a source temperature of 230 °C. Selected ion monitoring (SIM) mode was to monitor 77, 107, and 122 (m/z) ions with a gain factor of 10 and a scan speed of 1,562 (u/s).

### Phytoestrogen analyses

Similar methods to those used for 16S rRNA analyses were used in determining species differences in phytoestrogen analyte composition. Differences were visualized following nMDS of Bray-Curtis and Jaccard similarity indices, and following normalization, normality testing, and down-sampling, PERMANOVA was used to determine if species differences were observed (SWR, *n =* 16; GOHR, *n =* 16). Welch’s *t*-test was again used to measure significant difference between rhino species for individual analytes (SWR, *n =* 16; GOHR, *n =* 16). Since we did not observe a species-related difference using PERMANOVA, and no apparent clustering was observed with nMDS, hierarchical clustering (Bray-Curtis) was used to group phytoestrogen data into three profiles (A, B, C) based on their compositional similarity for further analyses. SIMPER analysis was used to determine contributions of each analyte to differences observed, and significant differences between groups were tested using analysis of variance (ANOVA; Profiles A, *n* = 23, B, *n* = 26, C, *n*= 9) with FDR correction.

### Receptor Activation

The ability of phytoestrogens and metabolites to activate SWR ERs was assessed using a SWR and GOHR estrogen receptor (ER) activation assay described previously by Tubbs *et al*. (2012; 2016) with minor modification. For each species, ER*α* or ER*β* sub-cloned into pcDNA3.1(+) expression plasmid (Invitrogen) was co-transfected into human embryonic kidney (HEK293) cells along with pCMX-β-galactosidase (β-gal), pGL2–3xERE luciferase reporter plasmid. After 24 hr cells were treated with phytoestrogens or metabolites, alone or in combination, and incubated for an additional 24 hr. For single test compounds, cells were treated with 100 pM-10 μM of each compound or vehicle (DMSO) alone. To assess the estrogenicity of phytoestrogen/metabolite profiles produced by SWR and GOHR microbial communities, cells were treated with serial dilutions of mixtures created *in vitro* to reflect those generated *in vivo*. Within each assay, a series of cells was treated with the endogenous estrogen, 17-estradiol (E_2_; 0.001-100 nM), to determine maximal E_2_ activation. Following incubation, cells were lysed and luciferase and β-Gal activity was measured as described previously (Tubbs *et al*., 2012). All data are presented as mean ± SE fold activation over vehicle treatment for each metabolite or mixture relative to the maximal activation E_2_. Differences in mean activation for both ERs were determined by ANOVA with FDR correction (each treatment, *n =* 9 (inter-assay, *n* = 3, intraassay, *n* = 3)), and significant interactions were observed between rhinoceros species and phytoestrogen profile. Data were sliced according the main effects, and differences within factor were observed.

### Fertility

Four calculations of SWR fertility were conducted for this study. Similar to Tubbs *et al*., (2016), two calculations are based on the number of offspring produced by the female per reproductive year through the completion of the study (Calf_study_; CS) and current levels (Calf_life_; CL). With pregnancies also considered a success, we conducted two additional calculations based on the number of pregnancies per reproductive year ending with the completion of the study (Pregnancy_study_; PS) through current (Pregnancy_life_; PL). GOHR samples were removed from the dataset so that data were not skewed by species differences.

### Correlations

Using the *microbiome* package in R (55), we examined significant correlations between OTUs (≥ 1.0 % relative abundance) and phytoestrogen analytes, OTUs and fertility measures, and those between phytoestrogen analytes and fertility measures. These correlations were carried out using the Spearman correlation method with multiple testing correction by FDR. OTUs found to correlate to fertility were further examined using a linear model (lm, *stats* package) (56).

## Acknowledgements

This work was supported by the Heller Foundation and the Ocelots Grants Program. The authors would like to thank San Diego Zoo Safari Park staff, Amanda Lussier and Jake Shepherd, and Rachel Felton (Institute for Conservation Research) for assistance in sample collection. We would also like to thank Mississippi State Chemical Laboratory staff, Ashli Brown, Darrell Sparks, Christina Childers, and Magan Green, for their assistance in mass spectrometry. Special thanks to Kimberly Dill-McFarland (University of British-Columbia) and Garret Suen (University of Wisconsin-Madison) for their aid in sequencing and their critical review of the manuscript. We would also like to thank all members of the Reproductive Sciences Team, Suen Lab, and Mississippi State Chemical Laboratory for their insightful discussions and support.

## Author contributions

CW and CT designed the experiment, and with AY and AM, CW conducted experimentation. Data analysis was carried out by CW. The manuscript was written by CW, CT, AM, and AY, with editorial assistance by BD.

